# Hypervariable-Locus Melting Typing (HLMT): a novel, fast and inexpensive sequencing-free approach to pathogen typing based on High Resolution Melting (HRM) analysis

**DOI:** 10.1101/2021.07.01.450706

**Authors:** Matteo Perini, Aurora Piazza, Simona Panelli, Stella Papaleo, Alessandro Alvaro, Francesca Vailati, Marta Corbella, Francesca Saluzzo, Floriana Gona, Daniele Castelli, Claudio Farina, Piero Marone, Daniela Maria Cirillo, Annalisa Cavallero, Gian Vincenzo Zuccotti, Francesco Comandatore

**Affiliations:** Department of Biomedical and Clinical Sciences “L. Sacco”, Pediatric Clinical Research Center “Romeo and Enrica Invernizzi”, Università Di Milano, 20157 - Milan (Italy); Department of Clinical, Surgical, Diagnostic and Pediatric Sciences, University of Pavia, 27100 - Pavia (Italy); Microbiology Institute, A.S.S.T. “Papa Giovanni XXIII”, 24127 - Bergamo (Italy); Microbiology and Virology Unit, Fondazione IRCCS Policlinico San Matteo, 27100 - Pavia (Italy); Emerging Bacterial Pathogens Unit, Division of Immunology, Transplantation and Infectious Diseases, IRCCS San Raffaele Scientific Institute, 20132 - Milan (Italy); Laboratory Microbiology and Virology -Ospedale San Raffaele Dibit, 2-San Gabriele 1, 20132 - Milan (Italy); Laboratory of Microbiology, ASST Monza, San Gerardo Hospital, 20900 - Monza (Italy); Department of Pediatrics, Children’s Hospital Vittore Buzzi, Università Di Milano, 20154 - Milan (Italy)

**Author notes:** corresponding author, Via Giovanni Battista Grassi, 74, 20157 Milano.

**Keywords:** Microbiological surveillance, High Resolution Melting, Outbreak reconstruction, Low-middle income countries, Real-time surveillance

## Abstract

**Objectives:** Subspecies pathogen typing is a pivotal tool to detect the emergence of high-risk clones in hospital settings and to limit their spreading among patients. Unfortunately, the most used subspecies typing methods (i.e. Pulsed-field Gel Electrophoresis - PFGE, Multi-Locus Sequence Typing - MLST and Whole Genome Sequencing - WGS) are too expensive and time consuming to be suitable for real-time surveillance. Here we present Hypervariable-Locus Melting Typing (HLMT), a novel subspecies typing approach based on High Resolution Melting (HRM) analysis, which allows pathogen typing in a few hours and with ∼5 euros per sample.

**Methods:** HLMT types the strains by clustering them using melting temperatures (HLMT-clustering) and/or by assigning them to Melting Types (MTs) on the basis of a reference dataset (HLMT-assignment). We applied HLMT (clustering and typing) to 134 *Klebsiella pneumoniae* strains collected during outbreaks or surveillance programs in four hospitals. Then, we compared HLMT typing results to PFGE, MLST and WGS.

**Results:** HLMT-clustering distinguishes most of the *K. pneumoniae* high-risk clones with a sensitivity comparable to PFGE and MLST. It also drawed surveillance epidemiological curves comparable to those obtained by MLST, PFGE and WGS typing. Furthermore, the results obtained by HLMT-assignment were coherent to MLST for 96% of the typed strains with a Jaccard index of 0.912.

**Conclusions:** HLMT is a fast and scalable method for pathogen typing, suitable for real-time hospital microbiological surveillance. HLMT is also inexpensive and thus it is applicable to infection control programs in low-middle income countries.

## Introduction

Healthcare-associated infections (HAIs) are a major burden for global public health [1]. The microbiological surveillance programs are pivotal to establish effective infection control strategies. In particular, subspecies typing is fundamental to detect the emergence of high-risk clones. The most used methods for subspecies bacterial typing are Pulsed Field Gel Electrophoresis (PFGE), Multi-Locus Sequence Typing (MLST) and Whole Genome Sequencing (WGS). All these methods require several hours (up to days) to be performed and this limits their application in nosocomial real-time surveillance programs.

High Resolution Melting (HRM) assay has been proposed as a suitable method for fast bacterial typing [2,3]. This technique measures the melting temperatures of qPCR amplicons, which depends on the GC content. HRM can even distinguish amplicons diverging for just one Single Nucleotide Polymorphism (SNP) and it is widely used to detect human allele variants [4]. HRM protocols designed on hypervariable genes are able to discriminate among bacterial clones within the same species [2,3], because their amplicons will melt at different temperatures (e.g. they differ in GC content). HRM is particularly promising for microbiological surveillance: it is fast (∼5 hours to complete the analysis), discriminatory, inexpensive (∼5 euros per sample) and it can be performed on the most common qPCR platforms [5].

Despite the numerous HRM protocols proposed so far for bacterial typing [2], the method is rarely applied in hospital settings for microbiological surveillance. Indeed, most protocols have been designed to distinguish only among the few clones used for protocol development, including only a fraction of the entire genetic variability of the pathogen. Moreover, only a few algorithms and software are available to analyse HRM data for epidemiological purposes.

Recently, we developed a novel approach for HRM-based subspecies typing. We focused on hypervariable genes and implemented a tool (i.e. EasyPrimer) to facilitate HRM primers designing in this difficult context [3]. Additionally, we developed an algorithm for pathogen typing using HRM data [6]. We used this approach to develop an HRM-based typing protocol for *Klebsiella pneumoniae* [3] and then we validated the repeatability and portability of the method [5].

In this study we provide a comprehensive description of this novel approach that we named Hypervariable-Locus Melting Typing (HLMT). We applied HLMT in four nosocomial epidemiological investigations on *Klebsiella pneumoniae*, comparing its typing efficiency to PFGE, MLST and WGS.

## Methods

### Ethics Statement

This study uses bacterial isolates from human samples that were obtained as part of hospital routines. No extra human samples were obtained for this research. Therefore, informed consent (either written or verbal) was not required.

### Isolates datasets

Four Italian hospitals were included in the study: “San Gerardo” Hospital in Monza (from here HSG), “IRCCS Fondazione Policlinico San Matteo” Hospital in Pavia (PSM), “ASST Papa Giovanni XXIII” Hospital in Bergamo (PG23) and “IRCCS San Raffaele” Hospital in Milan (OSR). The hospitals provided a total of 134 *Klebsiella pneumoniae* strains: i) HSG provided 10 strains isolated during an outbreak that involved 10 patients in the oncohematology ward between August 14, 2018 and September 24, 2018 (HSG dataset); ii) PSM provided 24 strains isolated during an outbreak already investigated with Whole Genome Sequencing (WGS) and described by Ferrari and colleagues [7] (PSM dataset) (the original dataset consisted of 32 strains, but only 24 were successfully revitalized in this work); iii) PG23 hospital provided 20 strains isolated during an outbreak that involved 16 patients and nine hospital wards, between May 07, 2019 and November 04, 2019 (PG23 dataset); iv) OSR hospital provided all the 80 strains isolated during a one-year-long WGS surveillance in 2017 and already typed with WGS and PFGE by Gona and colleagues [8] (OSR dataset).

### DNA extraction

For each of the 134 strains, the bacterial culture was subjected to two consecutive single colony selections on MacConkey agar, incubated overnight at 37 °C (Becton Dickinson, Franklin Lakes, NJ, USA). A single bacterial colony was then suspended in liquid medium, incubated overnight and the DNA was extracted using the DNeasy blood and tissue kit, following the manufacturer’s instructions (Qiagen, Hilden, Germany).

### Whole Genome Sequencing

The 104 *K. pneumoniae* strains isolated from PSM (n=24) and OSR (n=80) have been already subjected to WGS in previous studies [7,8]. The remaining 30 isolates (HSG and PG23 datasets) were subjected to WGS on the Illumina MiSeq platform, (Illumina, San Diego, CA, USA), after Nextera XT 2×250 bp paired-end library preparation. The reads were quality-checked using FastQC and trimmed using Trimmomatic software [9]. SPAdes[10] was then used to assembly the pair-end reads.

### WGS-based typing

The 15,699 public genome assemblies of *K. pneumoniae* present in the PATRIC database on February 8, 2021 for which the publication code was available (in accordance with Fort Lauderdale and Toronto agreements) were retrieved. Each of the four datasets was separately subjected to core SNPs calling as follows: i) each genome was compared to the retrieved PATRIC dataset using Mash [11] and the 50 most similar strains were included in the background dataset; ii) these selected PATRIC genomes were merged to the dataset genomes; iii) core SNP calling was performed on the merged genome dataset using Purple tool [8]. For each of the four datasets, the obtained core SNPs alignment was subjected to Maximum Likelihood (ML) phylogenetic analysis with 100 bootstraps using the software RAxML8 [12], after best model selection using ModelTest-NG [13] (GTR+G for HSG and PG23; TVM+G for PSM and OSR). For each of the four datasets, clusters were identified on the resulting trees as the largest monophyla of dataset strains (not from PATRIC) with a bootstrap support >= 75.

### Multi-Locus Sequence Typing and *wzi* alleles

*K. pneumoniae* clones are often defined by combining the MLST profile and the *wzi* gene allele [14]. Multi-Locus Sequence Typing (MLST) profiles and the *wzi* alleles of the 134 genome assemblies were determined using Kleborate [15].

### Pulsed Field Gel Electrophoresis Clustering

Pulsed Field Gel Electrophoresis (PFGE) clusters were described by Gona and colleagues [8] on the 80 strains of the OSR dataset following digestion with XbaI enzyme and separation into a CHEF-DRIII electrophoretic system (BioRad, Hercules, California).

### Hypervariable-Locus Melting Typing

HLMT includes two different typing strategies: HLMT-clustering and HLMT-assignment. HLMT-clustering groups the strains on the basis of melting temperatures without a reference dataset. HLMT-assignment classifies each strain into a Melting Type (MT) by comparing the melting temperatures of the strains to a reference dataset. The reference dataset used in this work was reconstructed using the 43 *K. pneumoniae* strains previously typed by HRM by Pasala and colleagues [5]: HLMT-clustering grouped the strains in seven clusters that we used to define seven Melting Types (MTs). For each MT, the reference melting temperatures were computed as the mid-range melting temperatures (the arithmetic mean of the highest and the lowest temperature) of the strains in that cluster. The MTs were then labelled using the names of the most relevant lineages of the strains that they contain. The reference dataset is available at https://skynet.unimi.it/wp-content/uploads/MeltingPlot/TemplateHLMT_ref_KPN_03-15-2021.xls. All the *K. pneumoniae* strains included in this work were subjected to High Resolution Melting (HRM) assays using the protocol described in Perini et al., 2020 [3]. For each of the four datasets, the obtained melting temperatures and the above-mentioned reference dataset were used to perform HLMT-clustering and HLMT-assignment analyses with MeltingPlot v2.0 tool [6] (available online at https://skynet.unimi.it/index.php/tools/MeltingPlot/).

### Comparison of the typing results

HLMT results were compared to PFGE, MLST+*wzi* and WGS by means of heatmaps and correlation plots, produced using the R libraries gplots [16] and corrplot [17], respectively. Furthermore, the HLMT-assignment results were compared to MLST+*wzi* by Jaccard similarity index, computed using the R package clustelval. For each dataset, HLMT-clustering, MLST+*wzi* and WGS typing results were combined to the collection dates of the strains to obtain the epidemiological curves showing the prevalence of the clusters and groups over time [6]. Additionally, PFGE typing results were retrieved from Gona et al. 2020 [8] and used to obtain the relative epidemiological curves.

### Data availability

Genome assemblies of the 30 strains sequenced in this work are available from the NCBI BioProject repository under project PRJEB44864 (ERP128959).

## Results

### Whole Genome Sequencing

The assembly statistics and the accession numbers of the 30 sequenced *K. pneumoniae* strains (10 of the HSG dataset and 20 of the PG23 dataset) are reported in the Supplementary Table S1.

### Typing methods results

Strains from the four datasets (HSG, PSM, PG23 and OSR) were typed by HLMT (clustering and assignment), MLST+*wzi* and WGS. Furthermore, the PFGE typing results of the 80 OSR dataset strains were retrieved from Gona and colleagues 2020 [8]. The results of HLMT, MLST+*wzi*, WGS and PFGE typing are reported in the Supplementary Table S1 and the core SNP-based phylogenetic trees used for WGS typing are shown in the Supplementary Figure S1.

### Comparison of the typing methods

For each dataset, the HLMT-clustering results were compared to MLST+*wzi* and WGS: the graphical representation of the relative contingency tables are reported in Supplementary Figure S2 and Supplementary Figure S3, respectively. HLMT-clustering results were compared to PFGE for the OSR dataset only (see Supplementary Figure S4). The HLMT-assignment algorithm classifies the strains into Melting Types. This analysis was able to classify 120 out of the 134 strains (∼90%) included in this study. Seven of these 120 strains (∼6%) belonged to MLST+*wzi* profiles not included in the HLMT reference strain dataset used for the analyses (see Methods). This made it impossible to assess, for these seven strains, if the HLMT-assignment and MLST+*wzi* results were coherent. For 108 out of the remaining 113 strains (96%) the HLMT-assignment and MLST+*wzi* were coherent, with a Jaccard similarity index of 0.912. The HLMT-assignment results are reported in the Supplementary Table S1 and the correlation matrix plot of the HLMT-assignment vs MLST+*wzi* is shown in Figure 1. For each dataset, the epidemiological curves obtained combining typing information (HLMT-clustering, MLST+*wzi*, WGS and PFGE) and isolation dates of the strains are shown in Figure 2.

**Figure 1:**
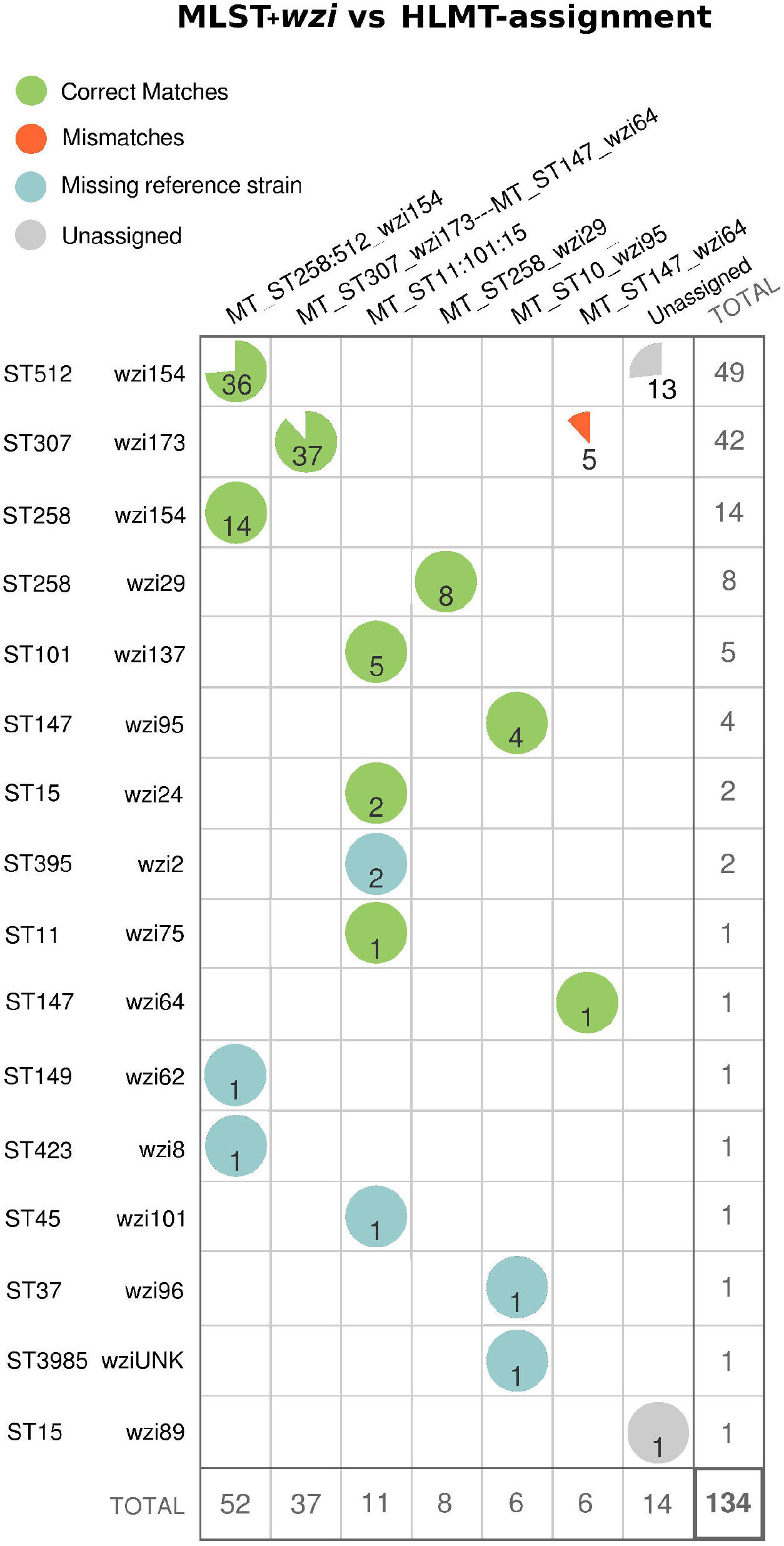
MLST+*wzi* vs HLMT-assignment correlation matrix. Correlation matrix between Multi-Locus Sequence Typing (MLST)+*wzi* typing and Hypervariable-Locus Melting Typing (HLMT)-assignment on the 134 strains analysed in the study. The pie charts indicate, for each MLST+*wzi* profile, the proportion of matches with the Melting Types (MT). Green pies show the correct matches and the red ones the mismatches. Light blue pies show the strains belonging to MLST+*wzi* profiles absent in the reference dataset used to perform HLMT-assignment analysis. Gray pies show the strains that were classified as “Unassigned” by HLMT-assignment analysis.

**Figure 2:**
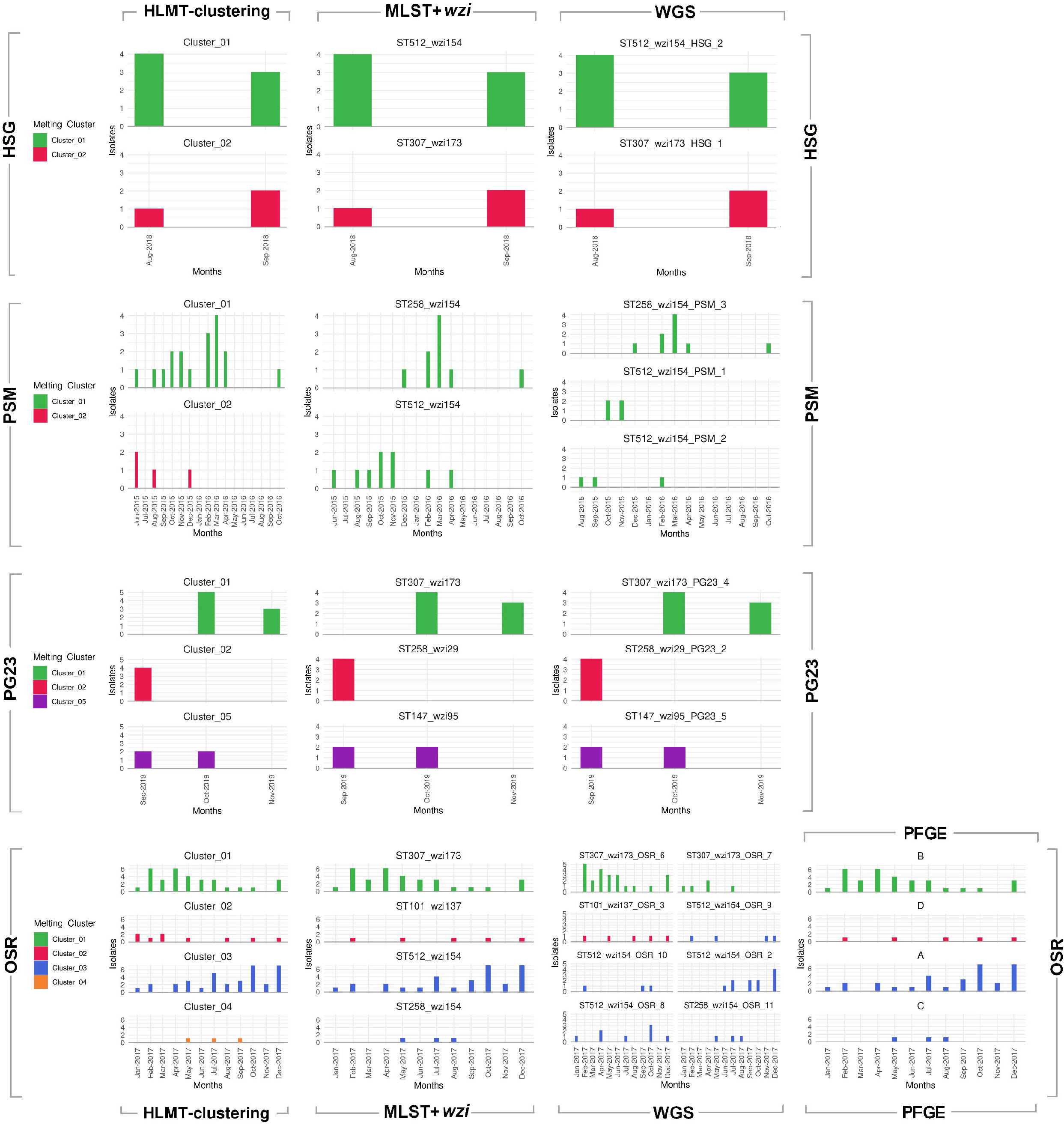
Epidemiological curves comparison. Epidemiological curves reconstructed using isolation dates and typing information obtained by HLMT-clustering, MLST+*wzi*, WGS and PFGE on the four dataset analysed in this study. Each line of the plot refers to a dataset while each column to a typing method. The colors used in the barplots correspond to the HLMT clusters.

## Discussion

Hypervariable-Locus Melting Typing (HLMT) is an innovative approach to High Resolution Melting (HRM)-based typing: it makes it easier to design highly discriminatory HRM protocols and to perform reliable, robust, repeatable and portable HRM-based typing analyses. In this work we show the application of HLMT to the typing of *Klebsiella pneumoniae* in four real hospital scenarios, comparing the results with those obtained by more established approaches, as Whole Genome Sequencing (WGS), Multi-Locus Sequence Typing (MLST) and Pulsed-Field Gel Electrophoresis (PFGE). As summarised in Figure 3, the workflow of HLMT consists of three main parts: i) HLMT protocol design on hypervariable genes; ii) HRM experiments; iii) HRM data analysis for HLMT-clustering and HLMT-assignment.

**Figure 3:**
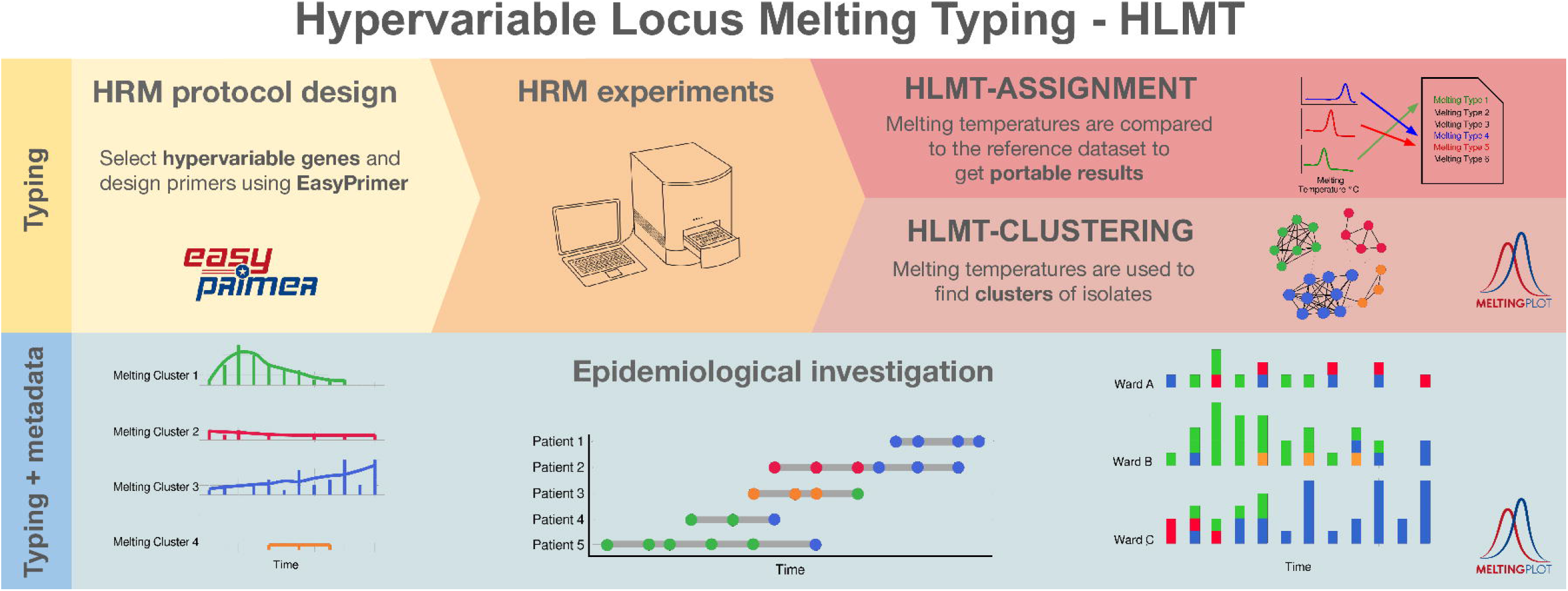
Hypervariable-Locus Melting Typing flow. Graphical representation of the flow of Hypervariable-Locus Melting Typing (HLMT). On the top, the three main steps of the HLMT method: HRM protocol design on hypervariable genes, HRM experiments and HRM data analysis for typing (HLMT-clustering and HLMT-assignment). On the bottom, the combination of HLMT typing and isolates metadata allow to perform epidemiological investigations, e.g. epidemiological curves and patient timeline. Primer design, HLMT-clustering, HLMT-assignment and epidemiological curve production can be performed using on-line free user-friendly tools (namely: EasyPrimer for primer design and MeltingPlot v2.0 for the others).

### HRM primer design

HRM is less sensitive than sequencing to discriminate among DNA sequences. Thus, to obtain a stronger signal, the HRM protocol should include more than one primer pair and it should be designed on hypervariable genes. Primer design on hypervariable genes is challenging because it requires the analysis of up to thousands of different gene alleles to identify conserved regions suitable for primer design. We already developed the EasyPrimer tool to automatically identify these gene regions [3].

### HRM experiments

HLMT analysis can be performed using HRM data obtained from any HRM-capable qPCR platform. As shown by Pasala et al. [5], HLMT-clustering is highly repeatable when the experiments are performed on the same model of instrument.

### HLMT-clustering and HLMT-assignment

HLMT analysis includes two different strain typing methods: HLMT-clustering and HLMT-assignment. HLMT-clustering groups the strains on the basis of their melting temperatures using a graph-based clustering algorithm [6]. Strains with similar melting temperatures for all the primers included in the HLMT protocol are grouped together. This clustering approach recalls the PFGE clustering, where a hierarchical clustering algorithm groups the strains on the basis of their restriction patterns.

On the other hand, in HLMT-assignment each strain is assigned to a Melting Type (MT) by comparing each strain to a reference dataset. The dataset consists of melting temperatures of previously typed strains selected to represent the genetic variability of the pathogen (see methods).

The strength of HLMT-clustering over HLMT-assignment is that the clustering works even on strains belonging to lineages absent in the HLMT strains dataset. On the other hand, the HLMT-assignment results are portable and they can be shared among laboratories and/or they can be compared to previous HLMT experiments.

The use of multiple primer pairs makes HLMT-clustering and HLMT-assignment more powerful but it also makes the analyses trickier. To tackle this issue, we have already developed MeltingPlot v2.0 tool [6] that automatically performs HLMT-clustering and HLMT-assignment analyses using HRM data.

### HLMT protocol for K. pneumoniae

We already designed an HLMT protocol for *K. pneumoniae* typing [3]. In this work, we show the applicability of the method for hospital surveillance programs. More in detail, we followed the three steps of the workflow described in Figure 3: i) HRM protocol design has been already carried out by Perini et al. 2020 [3], selecting the hypervariable capsular gene *wzi* as target and designing two primer sets (called *wzi*-3 and *wzi*-4) analysing the hundreds *wzi* allele sequences available, using EasyPrimer tool; ii) HRM experiments were performed in this work; iii) HLMT-clustering and HLMT-assignment analyses were also performed in this work using the HLMT reference strains dataset (see methods) and the melting temperatures of the 134 strains of the four datasets, using MeltingPlot v2.0 tool.

We reconstructed the *K. pneumoniae* reference dataset naming the Melting Types as the most epidemiologically relevant clone(s) present in the MT, recalling the labelling approach used in MLST for Clonal Clusters (e.g. *K. pneumoniae* CC258).

HLMT-assignment analysis classified almost every strain in concordance with MLST+*wzi* (Figure 1), showing the portability and repeatability power of the method. As shown in Figure 2, the HLMT-clustering epidemiological scenarios were highly consistent with those from the other typing methods: the curves are very similar for HSG, PG23 and OSR datasets and HLMT-clustering was also able to detect the emergence of the outbreak in PSM.

Furthermore, despite HLMT-clustering is less sensitive than MLST+*wzi*, the method was able to discriminate the most epidemiologically relevant STs (i.e. ST258 *wzi*29, ST258 *wzi*154, ST307 *wzi*173 and ST11) and the only not distinguished group of high-risk clones were ST101-ST11-ST15 and ST307-ST147 (Supplementary Figure S2). Here we also want to highlight that HLMT was able to discriminate between the two major sub-clones of the pandemic ST258, the Clade1 (harbouring the *wzi*29) and Clade2 (*wzi*154). These two clones, harbouring the same MLST gene alleles, are not distinguishable by MLST typing without further sequencing the gene *wzi*.

HLMT is less discriminatory than WGS but it is fast, easy, inexpensive and repeatable. The lower sensitivity of HRM in comparison to sequencing, makes HLMT unable to discriminate clones harbouring target genes with similar melting temperatures (e.g. ST101, ST11 and ST15 that harbour *wzi* allele with similar melting temperatures). Anyway, as shown in this work, this low sensitivity does not affect the repeatability of the method: indeed it fails to distinguish only some specific clones. On the other hand, this low sensitivity drastically reduces the possibility of obtaining false negative results. Therefore, when two strains are assigned to two distinct Melting Types (MT) or two Melting Clusters we can safely exclude their genetic relatedness. Having this information quickly during the early stage of a nosocomial outbreak can be particularly useful to establish an effective infection control strategy.

### Applicability

The only two instruments required to perform HLMT are a HRM-capable qPCR platform and a standard Personal Computer (PC); the molecular biology skills required are also minimal. No bioinformatic skills are necessary because both HLMT-clustering and HLMT-assignment can be automatically performed online, using the free and user-friendly tool MeltingPlot [6].

When the metadata of the strains is available, MeltingPlot can be used to merge HLMT clusters and strains metadata (isolation date and location) to produce graphical descriptions of the epidemiological scenario under study. Lastly, HLMT allows to type large amounts of isolates in a few hours for a few euros per sample (Figure 4).

**Figure 4:**
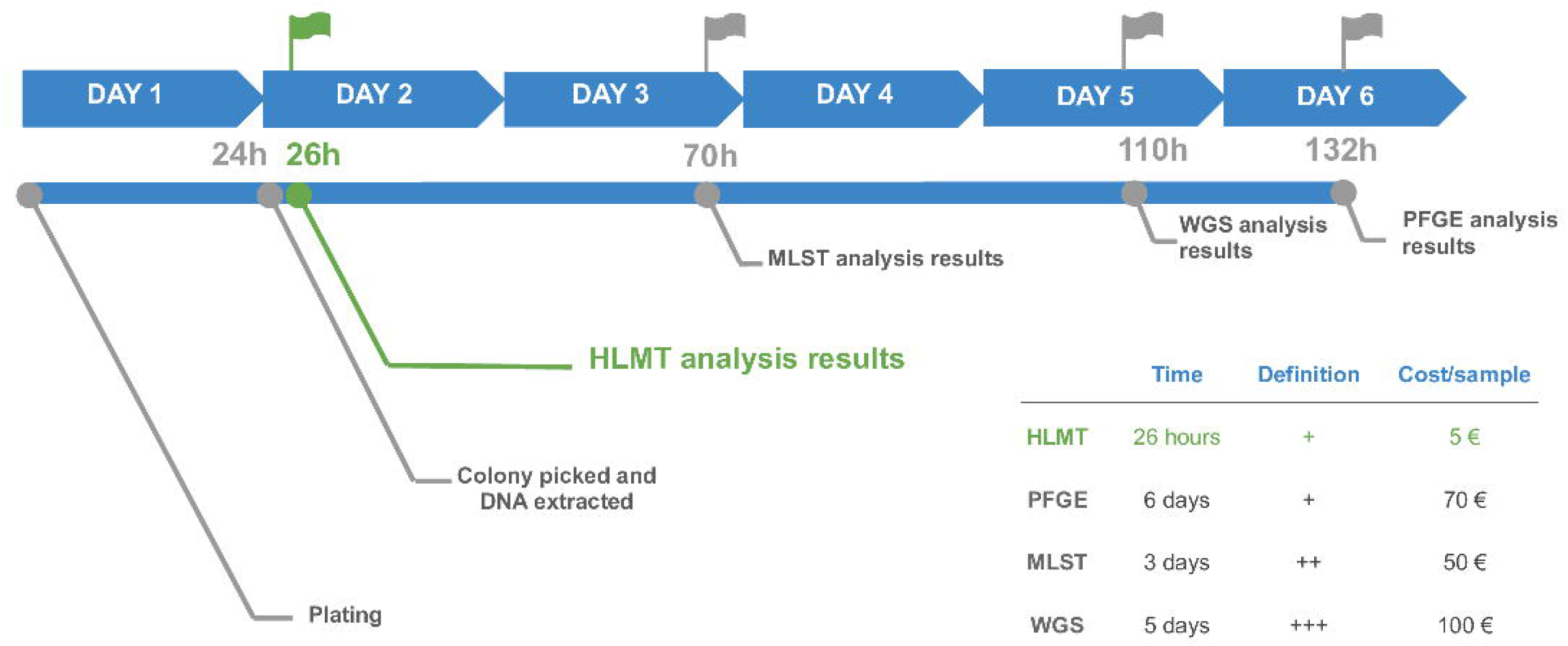
Timeline and costs of the typing methods. The timeline summarises the time required and the prices of Hypervariable-Locus Melting Typing (HLMT) (in green), Hypervariable-Locus Melting Typing (MLST), Whole Genome Sequencing (WGS) and Pulsed-Field Gel Electrophoresis (PFGE). The time required to perform each typing analysis, the relative typing definition and cost per sample are also reported.

WGS is the most sensitive method for bacterial typing and, even if it is becoming the gold standard typing approach for several pathogen species, PFGE and MLST are applied more often in real-time hospital surveillance programs [18]. In this work we show that HLMT is four times faster than MLST and 33 times faster than PFGE, maintaining a sensitivity comparable to these typing methods. Moreover, the low cost of HLMT (∼5 euros/isolate) makes it one of the most inexpensive sub-species pathogen typing methods available, being up to 20 times less expensive than MLST, PFGE and WGS (Figure 4). This makes HLMT suitable for epidemiological investigations in real-time and for the development of low cost surveillance programs even in low and middle income countries.

## Acknowledgements

Thanks to the Romeo ed Enrica Invernizzi Foundation.

## Supplementary Figures Captions

**FigureS1: WGS phylogenetic analysis**

Maximum Likelihood phylogenetic trees of the four datasets used in this study (HSG, PSM, PG23, OSR). Each dataset was analysed with the closest genomes found in PATRIC online database. The WGS cluster of each genome used in this study is shown by the colors next to the strain name.

**FigureS2: Contingency tables of HLMT-clustering vs MLST+*wzi***

For each of the four datasets analysed in this study (HSG, PSM, PG23, OSR), the HLMT clusters are compared to the MLST+*wzi* groups with a contingency table. Each color represents a HLMT cluster. The gray color indicates the strains that the HLMT clustering algorithm classified as “Undetermined”.

**FigureS3: Contingency tables of HLMT-clustering vs WGS**

For each of the four datasets analysed in this study (HSG, PSM, PG23, OSR), the HLMT clusters are compared to the WGS clusters with a contingency table. Each color represents a HLMT cluster. The gray color indicates the strains that the HLMT clustering algorithm classifies as “Undetermined”.

**FigureS4: Contingency table of HLMT-clustering vs PFGE**

For the OSR dataset, the HLMT clusters are compared to the PFGE clusters with a contingency table. Each color represents a HLMT cluster.

