## Supplementary figures and images for "Hypervariable-Locus Melting Typing (HLMT): a novel, fast and inexpensive sequencing-free approach to pathogen typing based on High Resolution Melting (HRM) analysis"

### Supplementary Figure S1

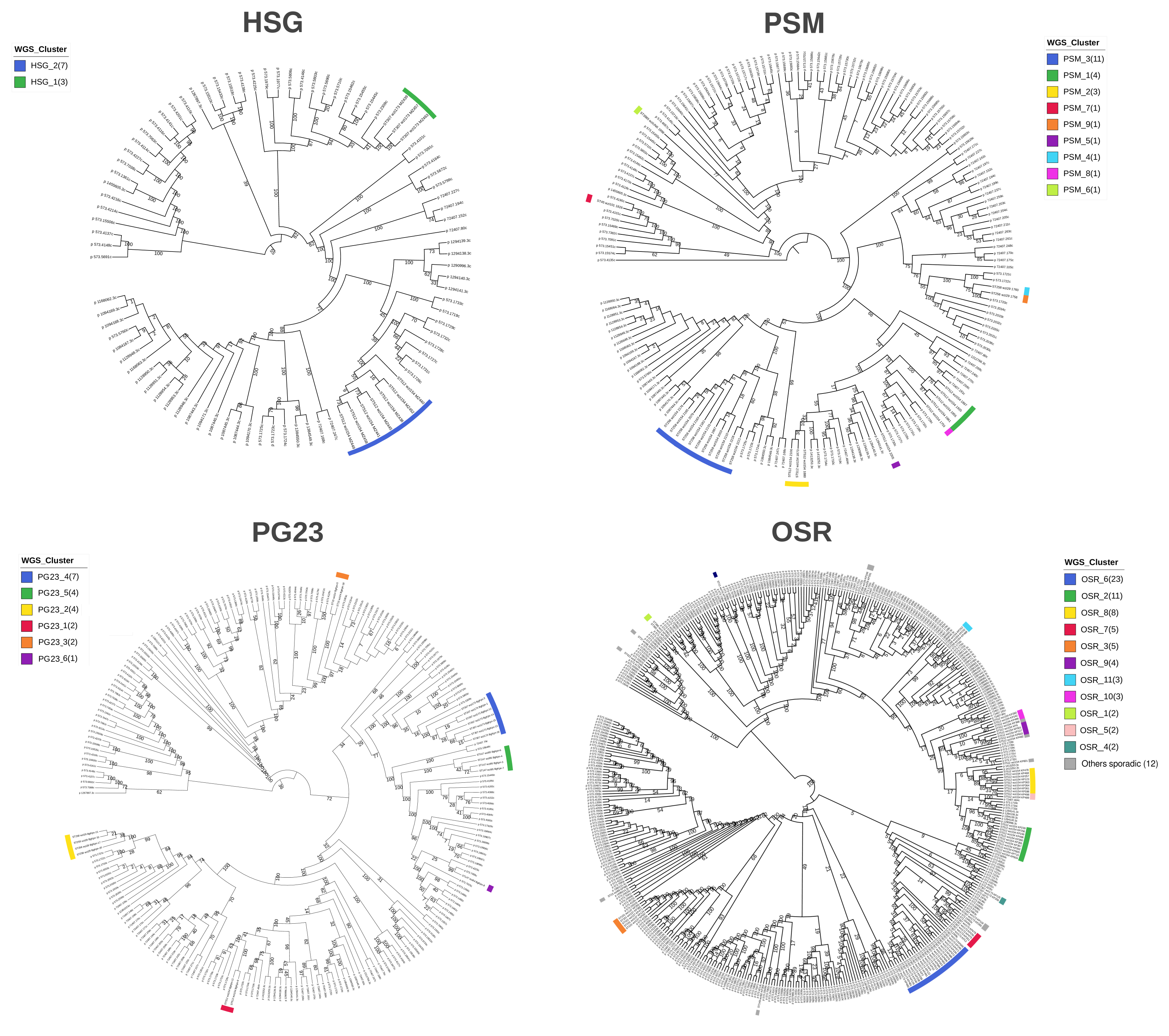

### Supplementary Figure S2

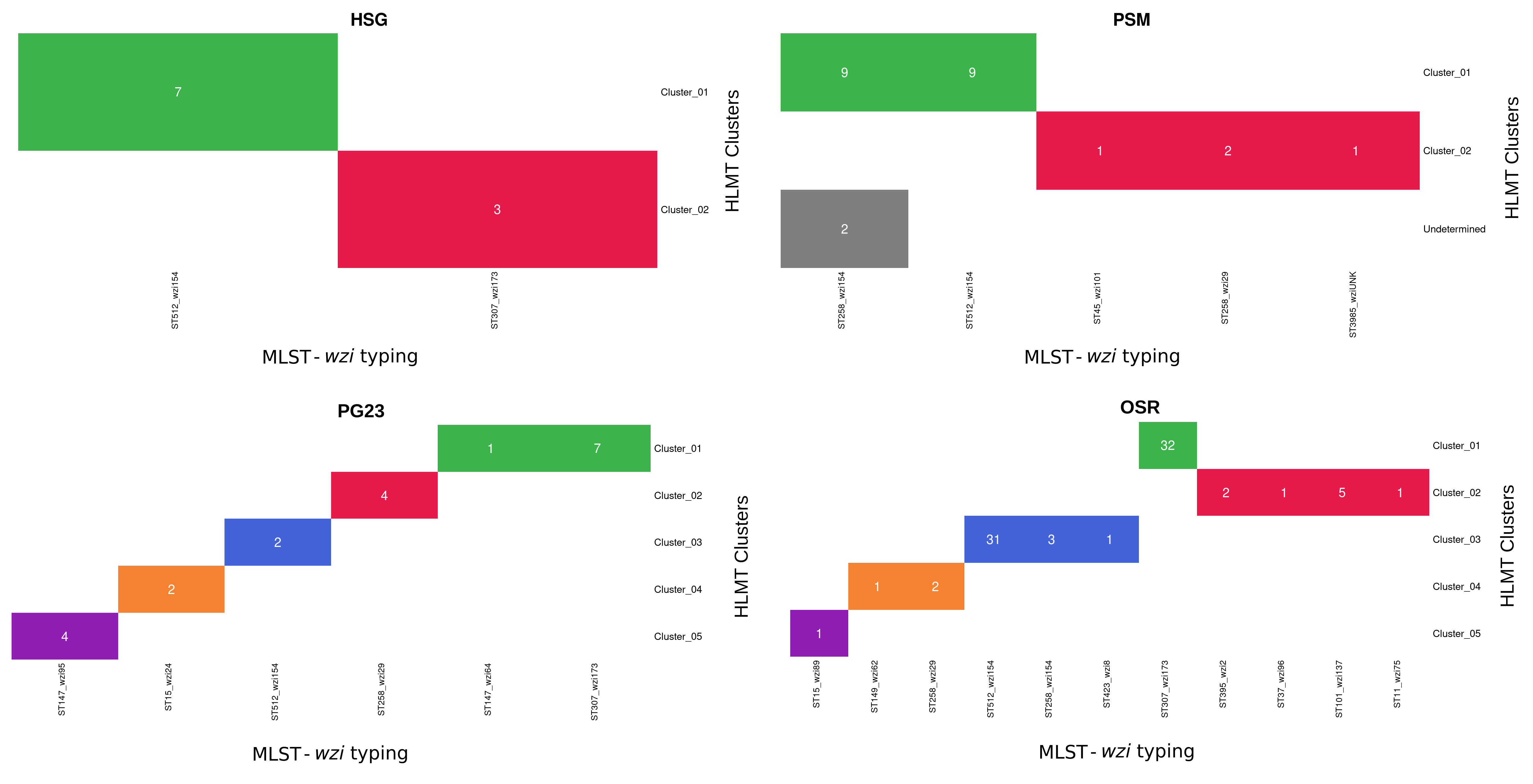

### Supplementary Figure S3

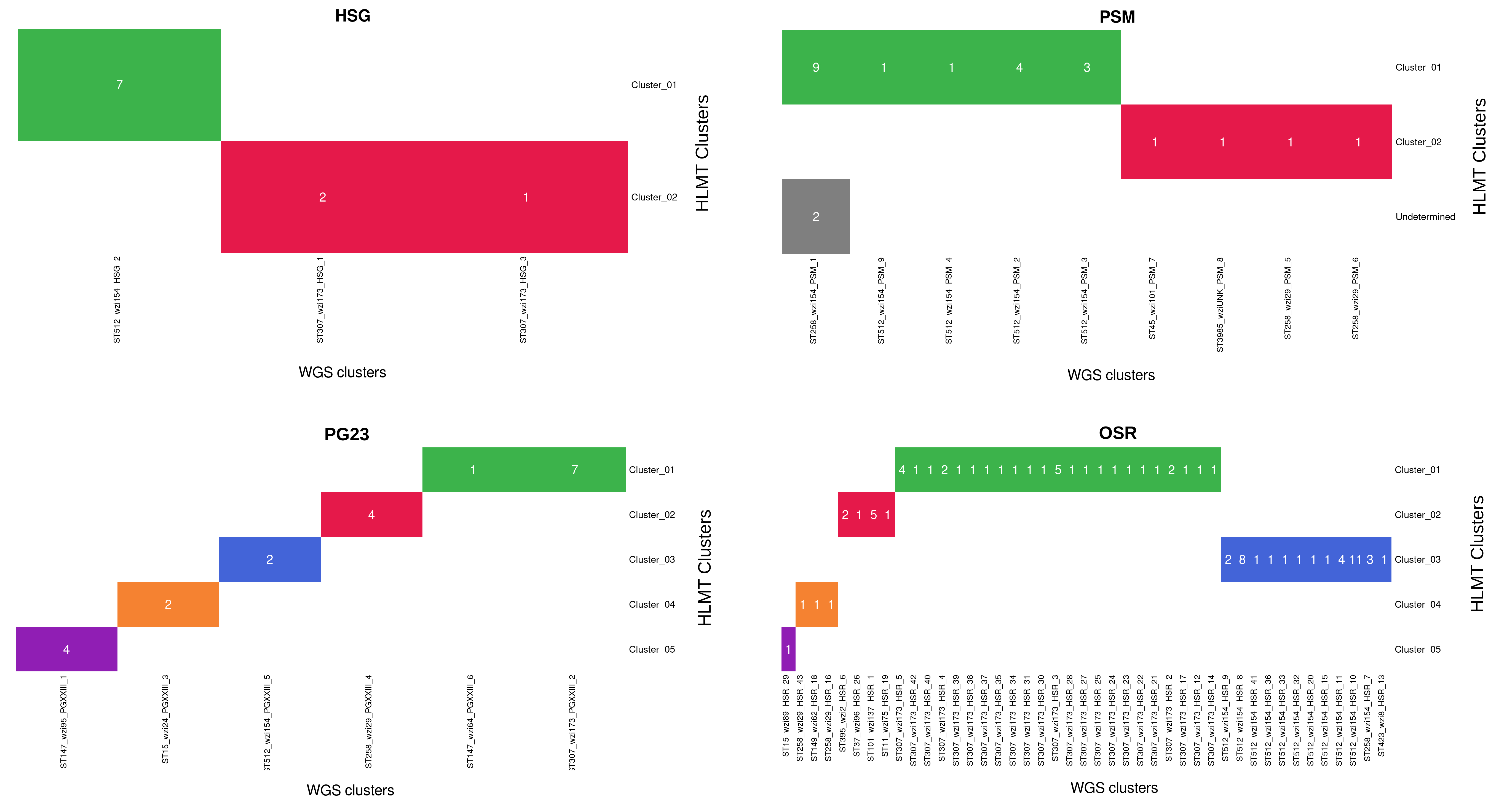

### Supplementary Figure S4

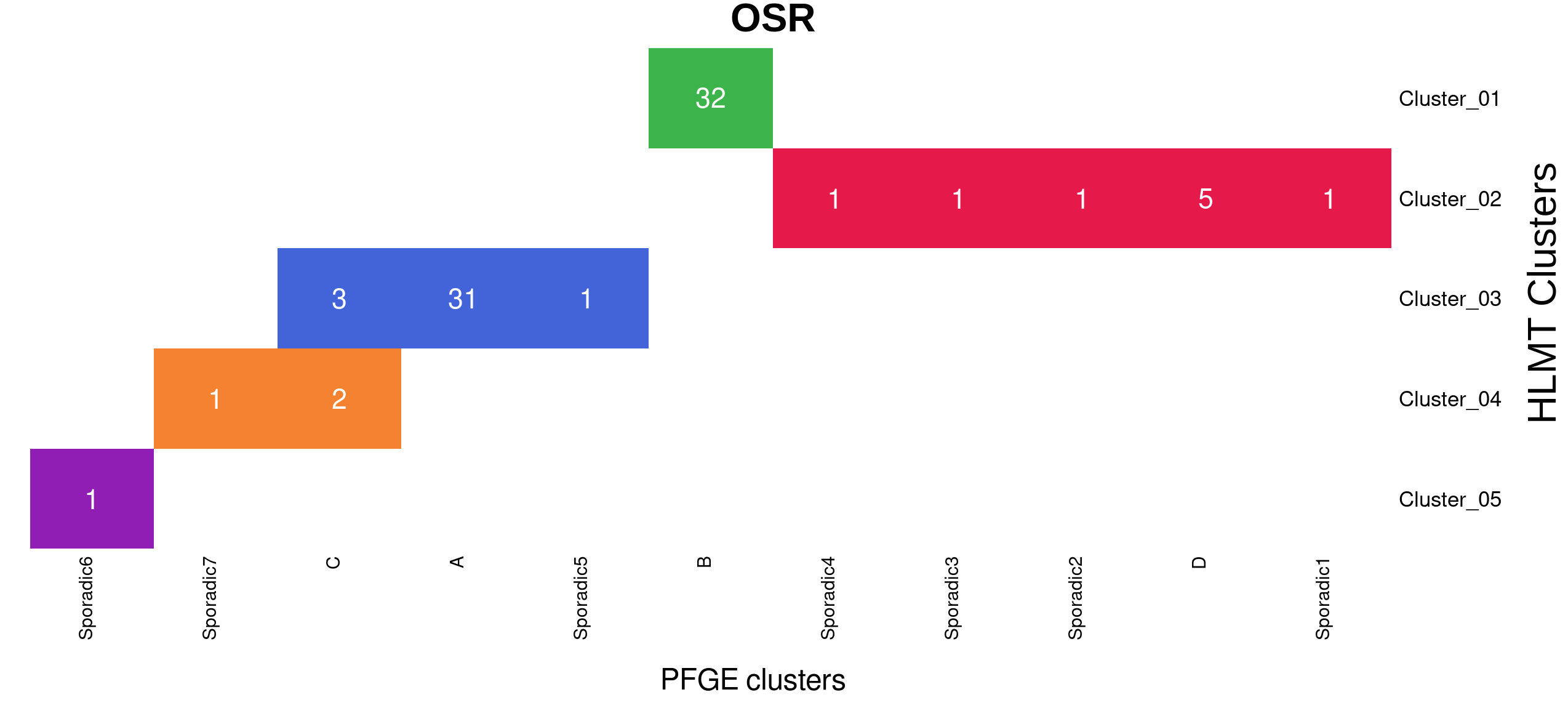
